# Stochasticity-induced stabilization weakens in diverse communities

**DOI:** 10.1101/2021.03.21.436309

**Authors:** Jayant Pande, Nadav M. Shnerb

## Abstract

Environmental stochasticity and the temporal variations of demographic rates associated with it are ubiquitous in nature. The ability of these fluctuations to stabilize a coexistence state of competing populations (sometimes known as the storage effect) is a counterintuitive feature that has aroused much interest. Here we consider the performance of environmental stochasticity as a stabilizer in diverse communities. We review the results of previous studies which suggest that the stabilizing effects of stochasticity weaken as the number of species increases, provide a systematic numerical exploration of the phenomenon and identify the relevant parameter regimes. Of particular importance is the ratio between the dwell time of the environment and the generation time: we show that stochasticity promotes diversity only when this ratio is smaller than the inverse of the fundamental biodiversity parameter. In an opposite regime, when stochasticity impedes coexistence and lowers the species richness, its effect is determined by the ratio between the strength of environmental variations and the rate at which new types are added to the community via speciation, mutation or immigration.

---

Coexistence of competing species poses a long-standing puzzle to the theory of community dynamics. The competitive exclusion principle states that the number of species is bounded by the number of limiting resources [1, 2], and May’s complexity-diversity analysis suggests that as the number of species increases their coexistence states become highly fragile [3]. Yet many natural communities, like fresh-water plankton [4], trees in tropical forests [5] and fish in coral reefs [6], seem to support high-diversity assemblages with very little evidence for substantial niche differentiation.

To resolve this problem, a few mechanisms that may facilitate coexistence have been presented and analyzed in the literature [7]. One of the most important has to do with the ability of stochastic temporal fluctuations to stabilize an otherwise unstable, or weakly stable, coexistence state [8, 9]. This counterintuitive phenomenon was discovered about forty years ago [10–12] and has been discussed since then by many authors. A few recent methodological papers have offered simple numerical methods to quantify the strength and the relative importance of this mechanism using empirical datasets [13, 14]. In parallel, several prominent studies have analyzed the role of environmental stochasticity in promoting the diversity of tropical forests [15, 16], grasses [17], yeast species [18] and *E. coli* strains [19]. The potential role of stochasticity-induced stabilization (SIS) in protecting genetic polymorphism has been considered as well [20–23].

Here we would like to discuss the relationship between the strength of SIS and the diversity of a given community. The basic observation is simple: all other things being equal, the ability of stochastic temporal variations to maintain coexistence *weakens* as the number of species grows. This weakening arises because the environmental effects on any given species get diluted, due to the differential response of all the other species to the environment. These weaker environmental effects, when superimposed on the destabilizing effect of demographic stochasticity at low densities, kill the impact of SIS. On the other hand, SIS strengthens as the dwell time of the environment (the time in which the environmental conditions stay more or less fixed) decreases. Therefore, stochasticity may increase diversity in rich assemblages only when the environmental fluctuations are rapid, i.e., if the correlation time of the environmental conditions is short.

In what follows we first review previous works from which these insights emerge, and then provide a systematic numerical exploration of the problem. Our numerical results suggest a quantitative criterion for the parameter regime in which SIS may promote diversity, namely that the dwell time of the environment must be smaller than the inverse of the fundamental biodiversity parameter. As the number of species grows, only rapidly changing environments may exhibit SIS.

These observations have a few interesting implications. First, the ability of SIS to support high-diversity assemblages turns out to be rather limited, unless the autocorrelation time of the environment is extremely short. Second, in some studies [15, 16] SIS for each pair of species has been quantified separately. Our work reveals the limitations of this methodology – the total effect of SIS in a diverse community is not necessarily proportional to its mean over species pairs. Finally, by considering a single population dynamics as a limiting case of a community with a large number of species, it becomes possible to understand why SIS never manifests itself in single population models (e.g., logistic growth with varying birth and death rates [24]).

## I. MODELS OF COMMUNITY DYNAMICS, WITH AND WITHOUT SIS

The general conditions under which stochasticity facilitates the coexistence of species are quite complex. Chesson and coworkers defined a set of scenarios under which environmental variations may stabilize biodiversity through the storage effect [8, 25]. These variations could be either stochastic or periodic in nature; the storage effect works in both cases [26–28]. Our main observation in this paper has to do with the buffering of environmental *stochasticity* in diverse communities, therefore we employ the term stochasticity-induced stabilization (SIS). While SIS is not identical to the storage effect, it overlaps with it [9], and in fact all our results here are demonstrated on versions of the lottery model which is the canonical example of a system exhibiting the storage effect [10, 11].

To provide a framework for our discussion and concrete mathematical models for the numerical analysis, we define here two individual based, continuous-time (Moran) versions of the lottery model. These versions have been considered in detail in [9, 29]. The continuous-time approach is more realistic than the discrete-time one, since it allows death and birth processes to take place in parallel, rather than requiring a clear separation between death periods and birth periods as in the original, discrete-time, lottery dynamics.

The use of continuous-time models allows us to disentangle the stabilizing and destabilizing effects of temporal environmental stochasticity. To do this, we will examine a local-competition model, in which the role of stochasticity is purely destabilizing (no SIS) [29–31], and a global-competition model, where stochasticity *may* stabilize coexistence.

- *Local competition* is perhaps the generic scenario for a community of animals who wander around in a certain spatial region, with an encounter between two individuals resulting in a struggle for some goods that affect survival and reproductive success, like a piece of food, a territory and so on. To model such an interaction we consider dynamics that take place via a series of duels between two randomly-picked individuals, in which the loser dies and the winner reproduces so the total number of individuals, *N*, is kept fixed. If the fitness of the first individual is exp(*s*_1_) and the fitness of the second is exp(*s*_2_), then the chance of the first to win the competition, *P*_1_, is determined by the fitness ratio

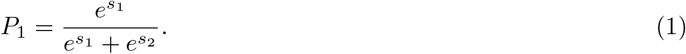 Analogously, *P*_2_ = 1 − *P*_1_.
- When the competition is global, one individual is chosen at random to die. The chance *P_a_* of any individual *a* to produce an offspring that recruits the resulting open gap is given by

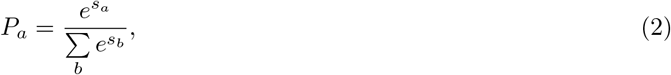

where the sum ranges over all the individuals. As a result, in the case when all the individuals of a species possess the same fitness, the chance of a given species *i* (with *n_i_* individuals) to capture the gap is

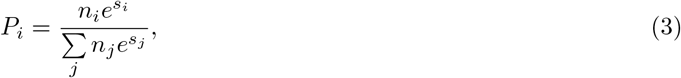

where the sum now ranges over all the species.

As mentioned, at any given instant of time all the individuals of a given species (say *i*) have the same log-fitness *s_i_*. In a temporally stochastic environment *s_i_* varies in time. In our numerical experiments, after each elementary birth-death event the chance of an environmental shift is 1/*Nδ*, so the dwell times of the environment (the durations of time during which the environment remains fixed) are picked at random from an exponential distribution whose mean is *δ* generations (with a generation being defined to be *N* elementary events). The autocorrelation time of the environment (i.e., the mean time until the next environmental shift, starting from a random point in time) is *δ*/2. After each environmental shift, a new and uncorrelated value for *s_i_* is picked for every species *i*.

Accordingly, at every instant of time each species *i* has the log-fitness *s_i_*(*t*) which is picked from a given probability distribution function *f_i_*(*s_i_*). If the mean of *f_i_*(*s_i_*) depends on *i*, then different species differ in their time-averaged fitness. In this case, in the absence of other stabilizing mechanisms one expects the (on-average) dominant species to select out all its competitors on a relatively short timescale. In contrast, when all the species have the same mean fitness, the mean time to extinction is much larger. Such “time-averaged neutral” dynamics [32, 33] provide the optimal conditions for SIS to manifest itself and to promote diversity, hence we consider these dynamics hereon. Accordingly, we assume that all the species draw their fitness from the same distribution *f* (*s*), which, for the sake of simplicity, has zero mean, so its main characteristic is its variance *σ*^2^. When *σ* = 0, both global and local dynamics are reduced to the standard (mainland, no spatial structure) neutral model that has been studied thoroughly in population genetics [34] and community ecology [35, 36].

In an individual-based model all species eventually go extinct and species richness reflects an extinction-speciation (or, in population genetics, extinction-mutation) equilibrium. In our simulations new types appear via the standard procedure of point speciation: at each birth, the offspring inherits the species identity of its parent with probability 1 − *ν*, and becomes the originator of a new species with probability *ν*. The response of a new species to environmental variations is independent of its parent species (i.e., it picks its *s*-value independently). In reality, the traits of a daughter type are correlated with the traits of the parent and one may thus expect a correlated response to environmental shifts. Here we simplify the picture by assuming that the parent and the daughter species are either completely correlated (in which case we consider them to be of the same type) or completely uncorrelated. The implications of this approximation are explained in the discussion section below. (Table I provides a glossary for the quantities considered through this paper.

**TABLE I.**
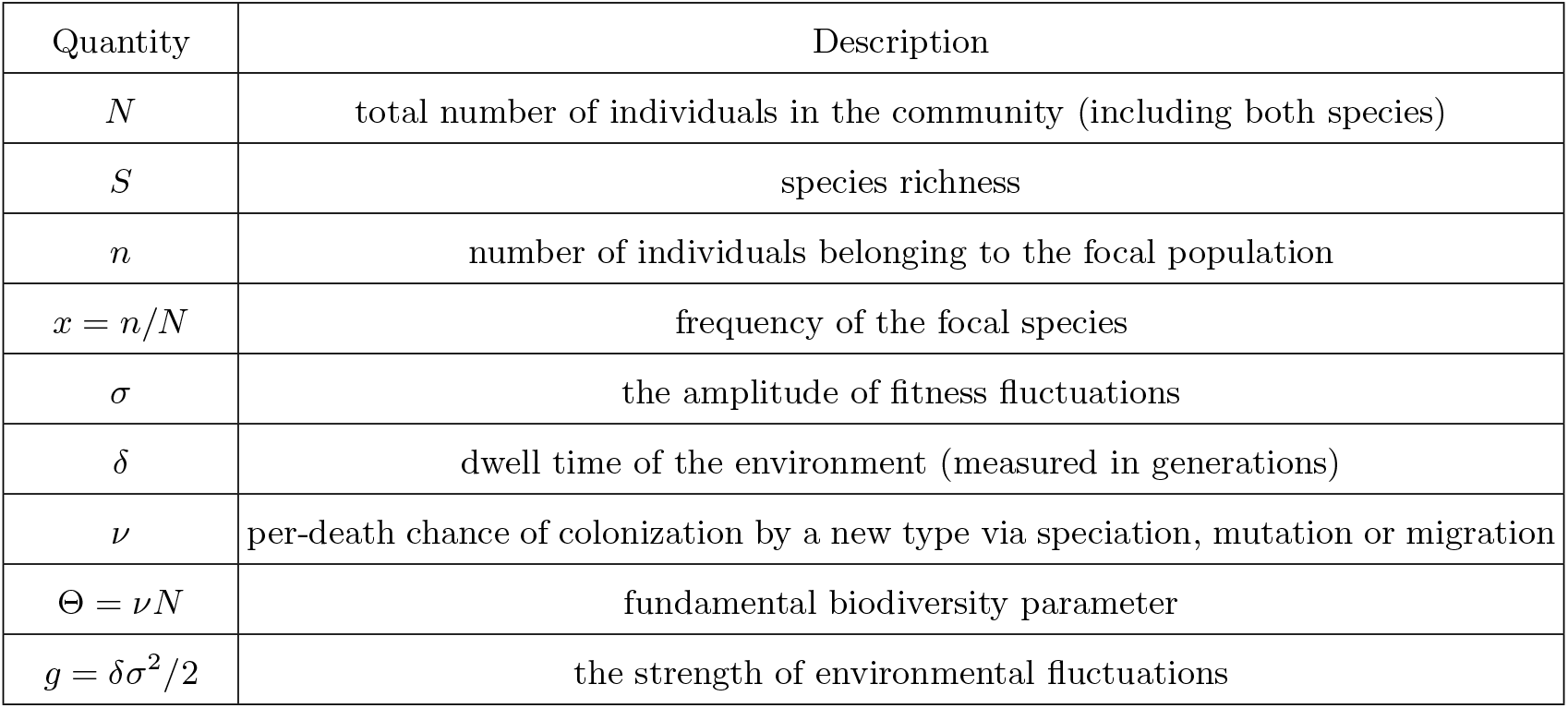
Glossary

To examine the stabilizing and the destabilizing effects of stochasticity, we compare the species richness at equilibrium with a baseline defined by the neutral model. As the strength of environmental fluctuations increases (i.e., as *σ* grows), one expects a decrease in the species richness in the absence of SIS, since abundance fluctuations grow and many species go extinct while the speciation rate is kept constant. Accordingly, with local competition an increase in *σ* decreases the species richness. In the presence of SIS the situation is more complicated. On one hand an increase in *σ* activates the mechanisms that stabilize coexistence, hence the number of species has a tendency to grow. On the other hand, these environmental variations still contribute to an increase in the abundance variations, which puts small populations in danger. Therefore, the main challenge is to understand the *σ*-dependence of the species richness in the global-competition model. A few theoretical insights that are relevant in this case are presented in the next section.

## II. REVIEW OF THEORETICAL INSIGHTS

In this section we first present the mean-field approach which simplifies the analysis in communities of many species and then we collect and recapitulate the most important theoretical findings of previous studies. All these studies, that employ different criteria for stability, point to the crucial importance of the dwell time. In the subsequent section III we present our numerical results which confirm this insight, clarify the relationship between dwell time and other characteristics of stochasticity like the amplitude of fitness variations, and identify the transition point beyond which stochasticity becomes a destabilizing factor.

### A. The mean-field approach

When the number of species is large, it is legitimate to adopt a mean-field approach, that is, to consider the dynamics of each individual population under the combined effect of all the other populations and the fluctuating environment. In our quantitative examples below we consider zero-sum dynamics, in which the total number of individuals summed over all the populations stays constant at *N*. This imposes some correlations among the abundances of the different species. Despite this, when the species richness *S* is large (*S* ≫ 1), these correlations are negligible and the mean-field approximation works quite well [33].

Use of this mean-field approach leads to the realization that the larger the species richness *S* is, the smaller is the amplitude of the environmentally-induced fitness variations (see [37] for a discussion of this “portfolio effect” in the context of intra-specific trait variability). Since *f* (*s*) is identical for all the species, the relative fitness of every single population reflects the difference between its own fitness (a random number, typically between −*σ* and +*σ*) and the mean fitness of an effective rival species, i.e., of an average individual in the rest of the community. The fitness of the effective rival population fluctuates through time, but since it reflects a sampling from *S* independent populations, its variance is smaller than *σ*^2^. When *N* is very large and *S* grows, the typical amplitude of fitness variations for the effective rival species is about 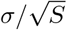 [33]. In other words, environmental effects are buffered in a diverse community, so while a given species in an *S* = 2 system is impacted by the fluctuations in its own fitness and in the fitness of its rival species, when *S* is large then every given species behaves as a single population in an inert (fixed-fitness) environment.

Interestingly, this argument alone, without any calculations, allows us to explain why the stabilizing effect of a fluctuating environment never manifests itself in single population models, like logistic growth with environmental stochasticity [24, 38–40]. This question is readily answered when one realizes that the dynamics of a single population are obtained as an effective field limit of the dynamics of a community with *S* → ∞. This is because in the latter case, from the perspective of any given population, the fitness of its effective rival species is fixed through time, and a species facing time-*independent* competition follows standard single-population logistic dynamics with environmentally-varying demographic rates. However, the mean abundance supported by SIS in a symmetric many-species system (i.e., one where all the species have the same mean fitness when averaged over time) equals *N*/*S*, and as *S* increases this corresponds to only a few individuals. Since demographic stochasticity dominates small populations [41], the stabilizing effect vanishes. When the model is asymmetric, then the situation is even worse, for then a given species is either inferior or superior to its effective rival and either disappears or takes over on short timescales.

### B. Models with fixed *S* and no demographic stochasticity the importance of the dwell time *δ*

In models with demographic stochasticity, a population is threatened by extinction if its abundance becomes low. Equivalently, even in models without demographic stochasticity, the stabilizing (or destabilizing) effect of stochasticity is measured by its ability to prevent populations from visiting the low-abundance region. Without demographic stochasticity, of course, populations cannot strictly go extinct so their properties have to be examined at fixed *S* (i.e., with no speciation and no extinction).

Two conditions are required to ensure that a population tends to avoid the low-abundance regime. First, the mean abundance supported by SIS (which in symmetric models is always *N*/*S*) has to be much larger than 1. Second, the single-species probability distribution function *P*(*x*), which is the chance to find a given species with *n = Nx* individuals (i.e., at frequency *x*), must decay close to *x* = 0. When *P*(*x*) diverges at *x* = 0 – even if the probability distribution is still normalizable – the system is less stable (as the introduction of demographic fluctuations would lead to extinction at small abundance populations [9, 11]).

A general solution for the joint distribution of species frequencies in the lottery model (with no speciation, no demographic stochasticity and no extinction) was presented by Hatfield and Chesson [12]. From their joint distribution one may extract a single-species distribution in the large-*S* limit, when the mean-field approximation is applicable and correlations among the abundances may be neglected [42]. In the time-averaged neutral case this yields

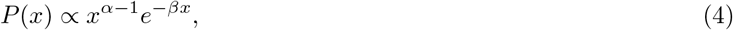

where *α* = (2/*S*)(1/*δ* − 1) and *β* = 2/*δ* − 3. This distribution diverges at 0 as long as *α <* 1, i.e., when *δ* > 2/(*S +* 2). This result suggests that the dwell time of the environment, *δ*, plays a major role in the stabilization of the system. SIS will facilitate coexistence at *S* ≫ 1 only if the dwell time of the environment is shorter than 2/*S* generations.

The results of Hatfield and Chesson [12] were obtained for a system without demographic stochasticity, using the diffusion approximation. In time-averaged neutral (symmetric) scenarios this case has a curious property: the outcomes (like *P*(*x*), and hence the estimated extinction times and so on) are independent of the amplitude *σ* of the environmental variations. As explained in [9, 42], this peculiarity has to do with the exact cancellation of the contribution of *σ* to stabilization (i.e., to the term in the equation that corresponds to deterministic growth or decay towards *N*/*S*) and its contribution to destabilization through stochastic abundance variations. Such a *σ*-independence of the results is clearly a problem of the theory – it appears to suggest that stochasticity-induced stabilization occurs even when the stochasticity vanishes – and to obtain results that make sense one must include demographic stochasticity in the model. Nevertheless, the main insight gathered from the Hatfield-Chesson solutions – the crucial role of *δ* and the need to decrease *δ* as *S* increases – turns out to be correct.

### C. Models with *S* = 2 and demographic stochasticity

When demographic stochasticity is included in a model and there is no supply of new types, an equilibrium state is a single-species state. The only remaining question is the persistence time of the diverse state: the larger this persistence time is, the more stable is the community.

The two-species lottery model, in which both the dwell time and the amplitude of fitness variations are explicit model parameters, was analyzed recently by Danino *et al.* [31]. It turns out that the scaling of the dwell time *δ* with the community size *N* affects the qualitative characteristics of the results. The case *Nδ* ≫ 1 may be considered as the generic scenario, since *δ* reflects the typical timescale over which environmental factors (such as temperature and precipitation) change. By and large, this scale has nothing to do with the size of the community, so as *N* increases the system always enters this regime. When *Nδ* ≫ 1 an increase in the amplitude of the environmental variations, *σ*, decreases the coexistence time in the local-competition model (which does not exhibit SIS) and increases the coexistence time in the global-competition model due to SIS. More qualitatively, while in the neutral case *σ* = 0 the persistence time (in generations) scales linearly with *N*, for local competition it scales like ln^2^ *N* (which is much shorter than *N*, so stochasticity destabilizes the community) while under global competition the persistence time scales like *N* ^1+1/*δ*^ (which is much higher than *N*, so stochasticity is a stabilizer).

A different kind of behavior is observed when *Nδ* ∼ 1 (i.e., when *δ* ∼ 1/*N*, which means that the environment flips after a small number of elementary birth-death events). For a local competition model without SIS the system converges to its neutral, *σ* = 0, limit. When competition is local there is no difference between the neutral case, where the chance to win a duel is 1/2, and the case where in each duel first the weather is picked at random and then the winner is picked using the probabilities for this weather: if the model is balanced, the net chance to win a duel is again 1/2. In contrast, in the global-competition case the stabilizing effect of stochasticity is strengthened as *δ* decreases, and it attains its maximal value when the chance of the environment to flip after each elementary step is 1/2, i.e., at *δ* = 2/*N*. More quantitatively, when *δ* ∼ 1/*N*, the persistence time of a two-species system grows *exponentially* with *N*, as opposed to the power-law growth when *Nδ* ≫ 1 [31].

Note that Adler and Drake [43] have considered a similar problem, namely the persistence time in a two-species model of annual plant dynamics with demographic stochasticity. However, in their system environmental variations govern only germination rates, and the effects of these rates on dwell times and fitness variations are quite complex. These authors observed a unimodal response of persistence times to the strength of these germination rate fluctuations, but an examination of the relationships between this fluctuation strength and our *δ* and *σ* requires more work.

### D. Models with demographic stochasticity and speciation

Under demographic stochasticity one may obtain a diverse community at equilibrium only due to the balance between extinction and the introduction of new types into the system via speciation, migration and so on. In this case the species richness is an emergent property of the system: *S* is not a parameter of the problem anymore, as it depends on the rate *ν* at which new types emerge and on the parameters that control the intrinsic dynamics of the community, i.e. the community size *N*, the strength of environmental fluctuations *σ* and the dwell time *δ*.

The single-species probability distribution function in that case (for both local and global competition, but only for *Nδ* ≫ 1) was calculated in [33]. Four main insights that emerge from this study are:

1. It is important to distinguish between “microscopic” and “macroscopic” species. Given *P*(*x*), the chance to find a population at a given frequency *x* = *n*/*N*, this distinction manifests itself in the nature of the decay of *P*(*x*) at large *x* values. When *P*(*x*) admits, say, an exponential cutoff at some *x*_c_ = *n*_c_/*N*, it implies that all species are microscopic, so their typical abundance is a fixed number that does not grow with *N* (unless *N* becomes very small, but we are interested here in large-*N* communities). In that case *S* grows linearly with *N* when *N* is large. On the other hand when *P*(*x*) decays slowly some species are macroscopic and a single population may compose a significant fraction of the whole community, say 1/10^th^ or 1/7^th^ of the total community size. In that case the growth of *S* with *N* is sub-linear. Therefore, diversity decreases when the system allows macroscopic populations.
2. A significant parameter is the ratio between *ν* and the effective strength of environmental stochasticity, *g* = *σ*^2^*δ*/2. Since the environment fluctuates, every species may encounter a series of years in which the environmental conditions favor its growth but it nevertheless suffers losses due to speciation that adds to its effective death rate. When 2*g*/*ν* is small, the losses dominate and all species are “microscopic”. In contrast, when 2*g*/*ν* is large environmental stochasticity dominates and the abundances of a few fortunate species are a finite fraction of *N*. Therefore, as 2*g*/*ν* increases the species richness decreases.
3. Accordingly, the parameter 2*g*/*ν* is the main scaling parameter of the problem. In models with local competition and no SIS the typical abundance of a population is fixed (*N*-independent) when 2*g*/*ν* is small. As *N* grows the number of different species with an independent response (*S*) grows and environmental effects are buffered. In this case the mean-field dynamics of any given species are approximately neutral. When 2*g*/*ν* is large the community supports some macroscopic populations at any given time and one expects a significant drop in the species richness with respect to the neutral case.
4. In the case of global competition, when the dynamics support SIS, the stabilizing effect adds an exponential cutoff on *P*(*x*) at 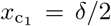 [see [33], Eq. (34)]. This implies that the species richness is larger than in the local-competition case, because no single species can grow too much and capture a substantial part of the community.

Based on point (4) above, one may suggest a simple way to determine the regime of parameters in which stochasticity may induce stabilization when it has the potential to do so (when the competition is global). In a neutral community with no spatial structure the chance of a species to have a given abundance follows the Fisher log-series [36, 44]. This log-series, Eq. (5) below, admits an exponential cutoff at 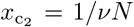, so species with frequency 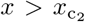 are rare. For SIS to increase the number of species, the cutoff it imposes, 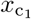, must be lower than that imposed by the neutral dynamics, i.e. 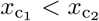, which implies *δ* < 2/*νN* or *δ*_c_ = 2/Θ, where Θ = *νN* is the fundamental biodiversity parameter. As we shall see, our numerical results suggest a critical value for *δ* which is about one-half of the prediction of this simple argument. Nevertheless, the argument yields the right functional dependence and the correct order of magnitude for the critical point.

## III. RESULTS

To confirm the general insights presented in the previous section, and to numerically characterize the important parameter limits that determine when stochasticity can and can not play a significant role in increasing the species richness of a community, we run Monte Carlo simulations of the local and global competition models described in section I and compare the results with the baseline defined by the well-mixed neutral model [36, 44, 45]. These simulations allow us to concretely identify the critical value *δ*_c_ beyond which SIS cannot increase the species richness. Additionally, our results demonstrate the universal nature of the species richness when seen as a function of the parameter 2*g*/*ν* (which the considerations in the previous section suggested was the main scaling parameter of the problem).

The species abundance distribution of the neutral model with the speciation rate *ν* is known to be given by the Fisher log series [36, 44, 45],

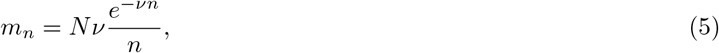

where *m_n_* is the expected number of species with abundance *n*. The total species richness in the neutral case is the sum over all *m_n_*, which gives

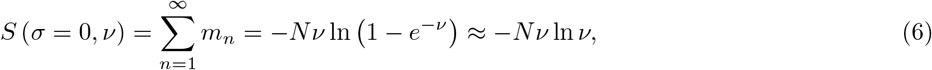

where the last approximation holds if *ν* ≪ 1.

Figure 1 shows the mean species richness as obtained from long, individual-based Monte Carlo simulations of our local-competition model with many values for the different parameters *N*, *σ* and *δ*. The species richness *S* is plotted against 2*g*/*ν*, the ratio between the strength of environmental variations, *g = σ*^2^*δ*/2, and the rate at which new types enter the system, *ν*. As expected, in this case environmental stochasticity always serves to lower the species richness below the *S*-value expected in the neutral case [Eq. (6)], since under local competition stochasticity only destabilizes. More remarkable is the data collapse: for all the many simulations with all the different values for the parameters *σ* and *δ*, once the species richness is plotted against 2*g*/*ν* the results collapse on to a single curve. As suggested in the previous section, stochasticity starts to affect the system when 2*g*/*ν* ≈ 1. When 2*g*/*ν* < 1 there are no macroscopic populations, the number of species grows linearly with *N*, and their differential response to the environment buffers the environmental stochasticity, which means that *S* is very close to its value expected in the neutral case. When 2*g*/*ν* > 1, at every instant of time some lucky populations that picked a rare series of good years constitute a finite fraction of the community, and the total diversity shrinks.

**FIG. 1.**
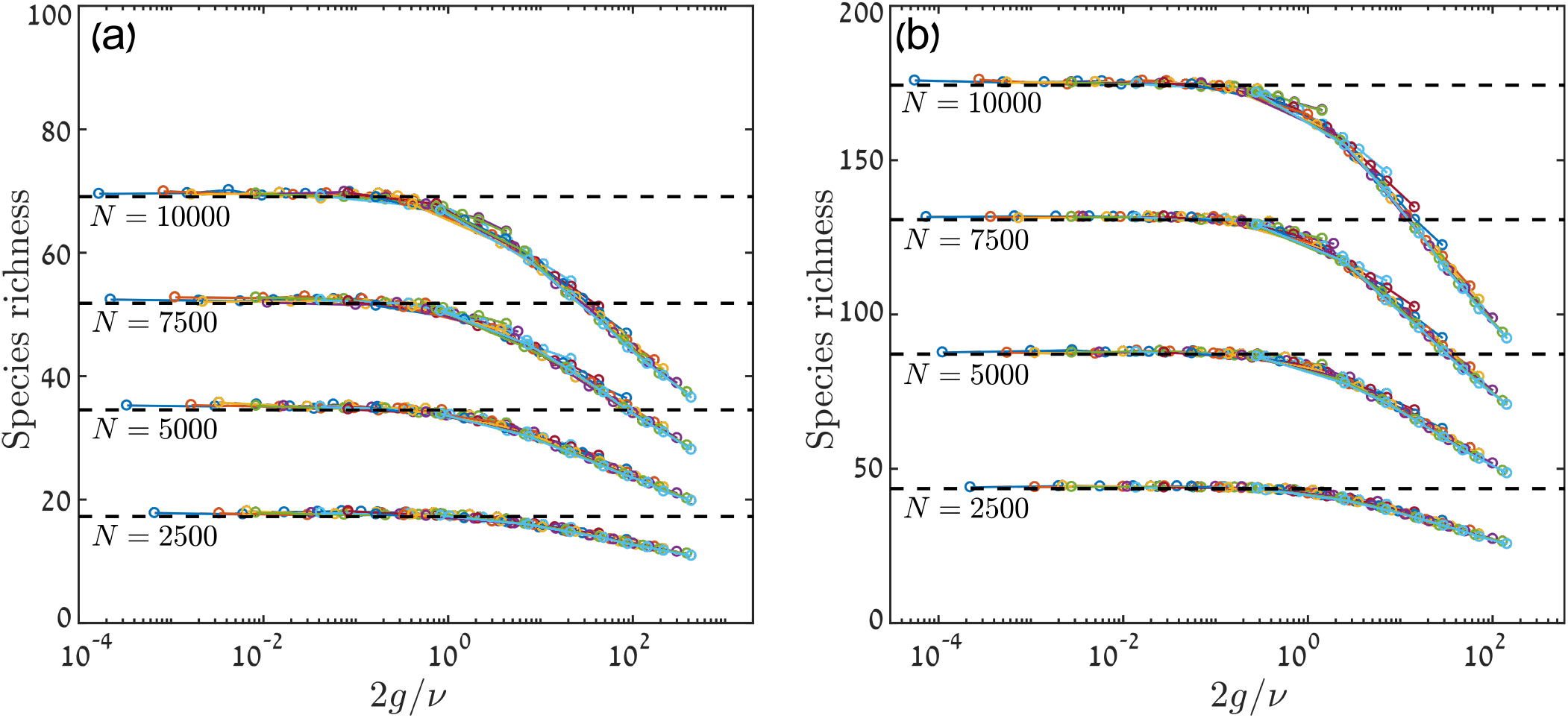
Species richness *S* as a function of 2*g*/*ν* for a model with local competition (i.e., without SIS). Results are shown for *ν* = 0.001 in panel (a) and for *ν* = 0.003 in panel (b). We ran long (10^5^ generations) simulations of the continuous-time (Moran) process with local competition, for four community sizes *N* = 2500, 5000, 7500 and 10000. In each of these sizes, we examined all possible combinations of thirteen values of *δ* (= 2/*N*, 10/*N*, 20/*N*, 100/*N*, 0.01, 0.05, 0.1, 0.2, 0.4, 0.5, 0.7, 0.9, 1.0) and ten values of *σ* (= 0.1, 0.2, 0.3, 0.4, 0.5, 0.6, 0.7, 0.8, 0.9, 1.0). Every run started with a homogenous population (i.e., a population with only one species), and was allowed to relax for 1000 generations. *S* was averaged over the history, from 1000 generations to the end of the run. Each color corresponds to a different value of *δ*, and different *σ*-values for each *δ* are connected with full lines. When *S* is plotted against the parameter 2*g*/*ν*, all the points (for a given value of *N*) collapse on to a universal curve. When 2*g*/*ν* ≪1, the results converge to the neutral limit (Eq. (6), dashed black lines). As 2*g*/*ν* grows, “macroscopic” species appear and the species richness declines.

Figure 2 presents the results for the case with global competition, i.e., when the dynamics support SIS. Now an increase in stochasticity may increase the species richness with respect to the neutral-model baseline. As expected, this happens only if *δ* is small enough. For each value of the community size *N*, above some critical value of the dwell time, *δ*_c_ (which depends on *N*), stochasticity becomes a destabilizer and an increase in *σ* leads to a decrease in *S*. When *δ* is even larger the results collapse on to a universal line when plotted against 2*g*/*ν*, as in the case of local competition. While the amplitude *σ* of the environmental variations plays an important role – when *σ* is small the system converges to its neutral limit, and the effect of environmental stochasticity on the species richness *S* clearly grows with *σ* – the *sign* of the effect depends only on *δ*.

**FIG. 2.**
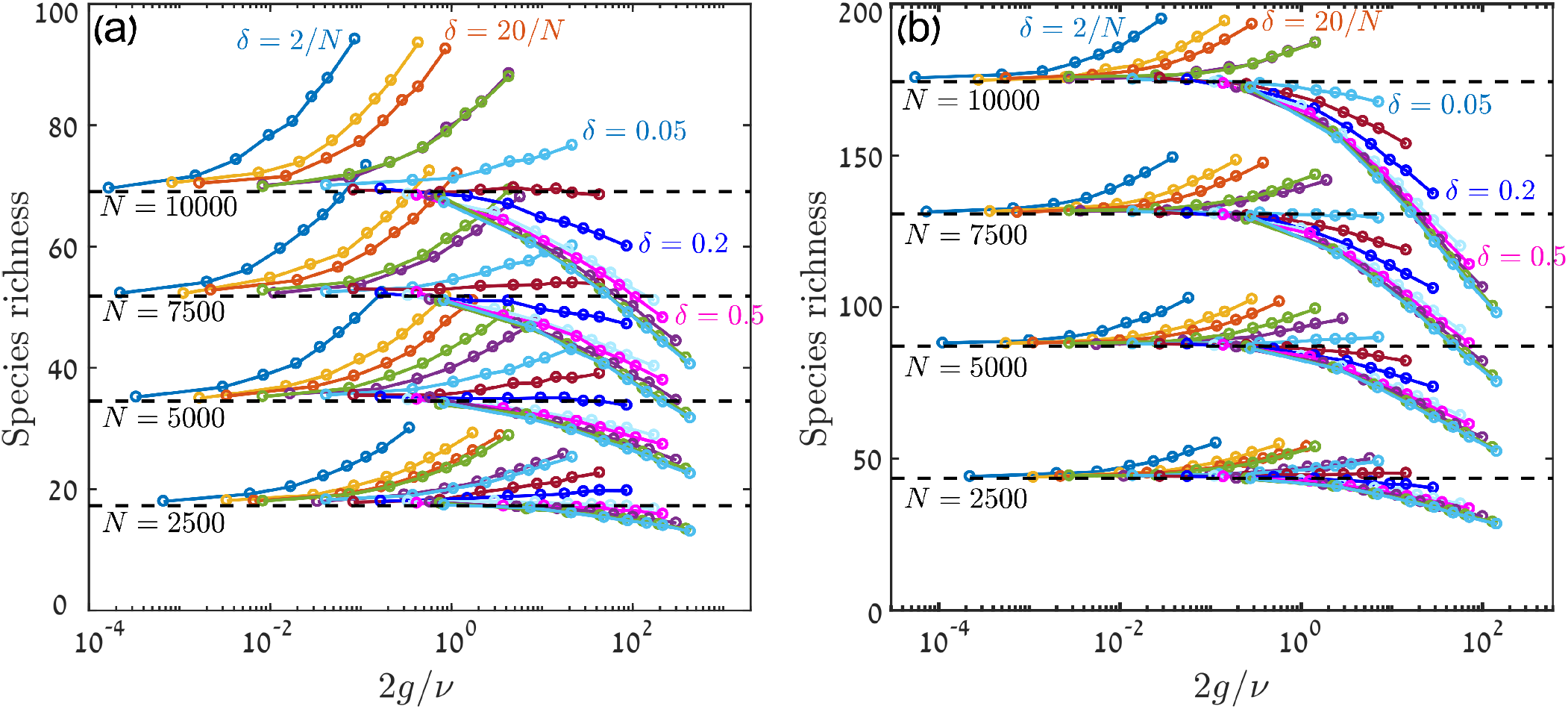
Species richness *S* as a function of 2*g*/*ν* for a model with global competition, in which stochasticity has the potential to facilitate coexistence. Results are shown for *ν* 0.001 in panel (a) and for *ν* = 0.003 in panel (b), with 10^5^ generations run in each simulation. Values of *N*, *σ* and *δ* are the same as used in Figure 1, namely *N* ∈ {2500, 5000, 7500, 10000}, *δ* ∈ {2/*N*, 10/*N*, 20/*N*, 100/*N*, 0.01, 0.05, 0.1, 0.2, 0.4, 0.5, 0.7, 0.9, 1.0}, and *σ* ∈ {0.1, 0.2, 0.3, 0.4, 0.5, 0.6, 0.7, 0.8, 0.9, 1.0}. Each color corresponds to a different value of *δ* – some of these *δ*-values are marked on the plots to indicate the general trend. Different *σ*-values for each *δ* are connected with full lines. As in Figure 1, the dashed black lines mark the results of the neutral case *σ* = 0 from Eq. (6). When *δ* is small, SIS leads to an increase of the species richness as *σ* increases. However, as *δ* increases the effect gradually disappears and above some critical value of *δ* (analyzed in Figure 3) stochasticity becomes a destabilizing factor and lowers the number of species. When *δ* increases even further, the data collapses on to a universal line and becomes (for a fixed *N*) a function of the single parameter 2*g*/*ν*.

We would like to stress that for a given set of parameters, *S* under global competition is always larger than *S* under local competition (to facilitate this comparison we used the same axes limits in Figures 1 and 2). In the regime *δ > δ*_c_ an increase in *σ* leads to a decrease in *S*, so this is not a regime where stochasticity induces stabilization, but even in this case when the competition is global the destabilizing effect of stochasticity is weaker.

What is the value of *δ*_c_? In the last section we presented a rough argument that suggests that *δ*_c_ = 2/*νN* = 2/Θ, where Θ = *νN* is the fundamental biodiversity parameter. We may now use our numerical results to provide a better estimate for this quantity. For *δ < δ*_c_, an increase in *σ* increases the species richness (due to SIS), and when *δ > δ*_c_, species richness decreases with *σ*, so at the critical value *δ*_c_ one expects a *σ*-independent species richness. As shown in Figure 3, the identification of this *δ*-value is straightforward, and our numerics suggest that *δ*_c_ = 1/Θ = 1/*νN*, half of the value predicted by our theoretical argument.

**FIG. 3.**
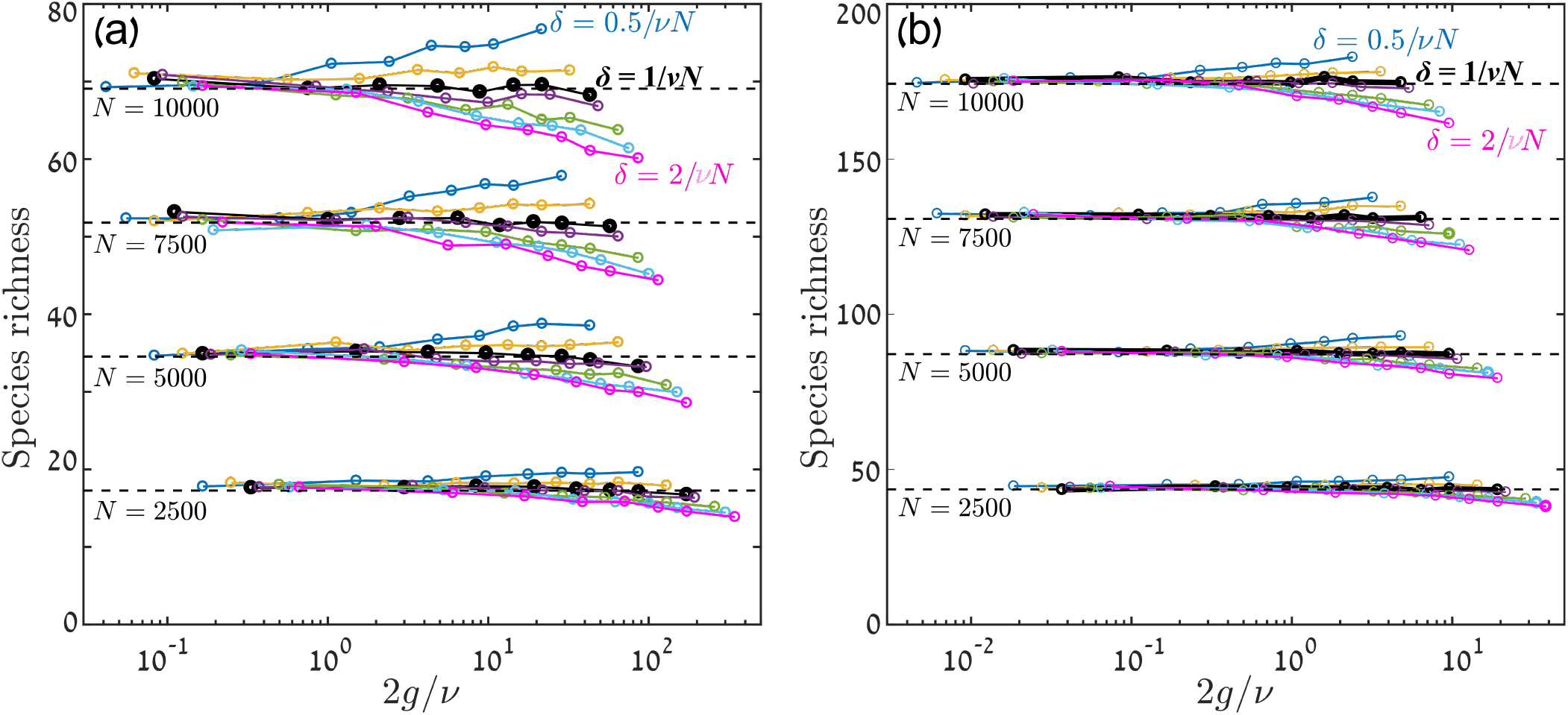
Species richness *S* as a function of 2*g*/*ν* for a model with global competition and SIS, for *δ*-values around the critical value. Results are shown for *ν* = 0.001 in panel (a) and for *ν* = 0.003 in panel (b), with 30000 generations run in each simulation. The values of *N* and *σ* are the same as used in Figures 1 and 2, namely *N* ∈ {2500, 5000, 7500, 10000} and *σ* ∈ {0.1, 0.2, 0.3, 0.4, 0.5, 0.6, 0.7, 0.8, 0.9, 1.0}. The *δ*-values used are *δ* ∈ {0.5/Θ, 0.75/Θ, 1.0/Θ, 1.125/Θ, 1.5/Θ, 1.75/Θ, 2.0/Θ} (where Θ = *νN*) and some of these values are marked on the plot. The dashed black lines mark the results of the neutral case from Eq. (6). The results suggest that the transition from a stabilizing to a destabilizing effect of stochasticity takes place at *δ*_c_ = 1/Θ (shown in thick black lines), where the species richness becomes *σ*-independent. At *δ*_c_ the system sticks to its neutral-model species richness even when *σ* is large.

## IV. DISCUSSION

All environments fluctuate, and their fluctuations affect the demographic rates of populations. When this effect is correlated among the different populations in a community, it only changes the effective population sizes, but the relative fitnesses of the different populations are fixed through time. In this case, if the value *N* of the total community size reaches a very low level (i.e. a bottleneck) one expects a significant loss of biodiversity, otherwise the effect on extinction times (and hence on the species richness) is relatively weak [46–49].

The situation changes when species demographic rates respond *differentially* to the environmental variations (in other words, when there are species-specific environmental responses). In this case there are two scenarios. When the dynamics do not support SIS, the only effect of these temporal stochastic fluctuations is to increase the abundance variations and to decrease the extinction times, hence the species richness decreases. As we have shown through our simulations above, the relevant scale is 2*g*/*ν*, where *g* is a measure of the strength of the environmental stochasticity and *ν* is the rate at which new types arrive in the community (due to processes like mutation, speciation and migration). When 2*g*/*ν* is smaller than 1 there is almost no effect of the changing environment and the species richness is very close to the one obtained for neutral dynamics in a fixed environment. When 2*g*/*ν* > 1 a few macroscopic species emerge (whose abundance is a finite fraction of *N* and grows linearly as *N* increases) and the species richness decreases.

When the system supports SIS, as in the case of global competition, then the effect of stochasticity may increase its species richness with respect to the neutral case, but for that to happen the dwell time of the underlying environmental fluctuations has to be below *δ*_c_, so it should be inversely proportional to the fundamental biodiversity parameter *νN*. Therefore, when all the other parameters are kept fixed and *N* increases, *δ*_c_ goes to zero and the effect of stochasticity becomes destabilizing, with the species richness falling below the neutral-case baseline.

In both the cases (with and without SIS) one may adopt the mean-field approach and consider the dynamics of a single species under the effect of all the other species that form together an effective rival population. As *N* → ∞ the species richness diverges, so the fluctuations in the fitness of the effective rival species (i.e., the fluctuations in the mean fitness of an individual that does not belong to the focal species) vanish. In this case the system becomes a collection of single populations, each interacting with a nearly-fixed effective environment. Stochasticity then becomes a purely destabilizing mechanism, the time to extinction shortens, and the species richness is lowered.

In real communities, some groups of species share important traits and respond to environmental changes with a degree of coherence [50, 51]. In general one expects these correlations to increase dramatically when *S* becomes larger than the number of available temporal niches [28]. In this work we assumed fully-independent responses of different species to environmental variations, so we neglect correlated response and allow an unlimited number of temporal niches. In order to apply our insights to real communities it would be necessary to identify our “species” with functional groups, such that above some coherence threshold, which would need to be defined, species would be considered as belonging to the same functional group. Correspondingly, what we call here the “speciation rate” *ν* would have to be replaced by the rate at which new functional groups (with uncorrelated responses to environmental variations) emerge during the dynamics. This latter rate is smaller than the rates inferred from standard taxonomic analyses.

Due to all this, the assessment of the role of stochasticity in stabilizing the coexistence of diverse assemblages is not a simple task. For example, Usinowicz and coworkers [16] measured the stabilizing effect of temporal stochastic variations for each *pair* of tree species in forest communities, with the mean of all these partial effects serving as a metric for the overall stabilizing effect. As we have seen, such an approach needs to be examined carefully. One should, first of all, study the correlations in the responses of different species to the environment and consider functional groups together. Even if the number of functional groups is large (for instance when taxonomic species indeed have primarily uncorrelated responses to the environment), the effect of environmental stochasticity is buffered as *S* increases and the dwell time of the environment must be very small for SIS to increase the species richness. If, on the other hand, the number of functional groups is small, then each of these groups likely contains many species with neutral competition between them. In such a case, SIS stabilizes not species but sub-communities (i.e. functional groups), with neutral dynamics within each sub-community. Since the number of species under neutral dynamics is proportional to the abundance of the sub-community, the contribution of SIS to the overall species richness is rather weak [16].

What then are the conditions under which SIS may contribute to coexistence in diverse communities? Our analysis suggests two possible routes. First, it may happen that, although the community is diverse, the number of effective functional groups that have a differential response to environmental variations is small, maybe 2 or 3. In that case the storage effect contributes to the coexistence of these groups, but other mechanisms must be invoked to explain within-group coexistence, so SIS may play a role in a hierarchy of coexistence mechanisms.

Another possibility is the case of very small *δ*. This may happen in populations with large generation times. For example, the lifetime of a tree in a typical tropical forest is about 50 years. If environmental variations are uncorrelated between years, as suggested in [15, 16], then *δ* ∼ 0.02, so *δ* may be smaller than *δ*_c_ even in large-*S* communities. Since the amplitude of variations *σ* plays a secondary role with respect to the autocorrelation time *δ*, it may happen that short-term, low-amplitude variations yield the required effect. Note that these variations must be correlated over space, otherwise the stochasticity affects conspecific individuals differently and this only increases the effective strength of demographic stochasticity.

All in all, we feel that the existing literature does not stress enough the dependence of SIS and of the storage effect on the diversity of a system. Our analysis emphasizes the need to assess a community on the whole [52], with special attention paid to interspecific correlated responses and to the buffering of the environmental variations when the response is differential. We hope that this work will motivate more informed studies of environmental stochasticity and its effects on species coexistence, leading to a better understanding of the forces that shape biodiversity in our world.

## V. ACKNOWLEDGMENTS

This research was supported by the ISF-NRF Singapore joint research program (grant number 2669/17).

